# An extension to: Systematic assessment of commercially available low-input miRNA library preparation kits

**DOI:** 10.1101/2020.01.25.919431

**Authors:** Fatima Heinicke, Xiangfu Zhong, Manuela Zucknick, Johannes Breidenbach, Arvind Y.M. Sundaram, Siri T. Flåm, Magnus Leithaug, Marianne Dalland, Simon Rayner, Benedicte A. Lie, Gregor D. Gilfillan

**Affiliations:** Department of Medical Genetics, Oslo University Hospital and University of Oslo, Oslo, 0450 Norway; Department of Biostatistics, Oslo Centre for Biostatistics and Epidemiology, University of Oslo, Oslo, Norway; Norwegian Institute for Bioeconomy Research, National Forest Inventory, Ås, 1431 Norway

**Keywords:** microRNA, miRNA, small RNA-seq, library preparation, sequencing bias, low RNA input, NGS, Next Generation Sequencing, NEBNext, NEXTflex

## Abstract

High-throughput sequencing has emerged as the favoured method to study microRNA (miRNA) expression, but biases introduced during library preparation have been reported. To assist researchers choose the most appropriate library preparation kit, we recently compared the performance of six commercially-available kits on synthetic miRNAs and human RNA, where library preparation was performed by the vendors. We hereby supplement this study with data from two further commonly used kits (NEBNext, NEXTflex) whose manufacturers initially declined to participate. As before, performance was assessed with respect to sensitivity, reliability, titration response and differential expression. Despite NEXTflex employing partially-randomised adapter sequences to minimise bias, we reaffirm that biases in miRNA abundance are kit-specific, complicating the comparison of miRNA datasets generated using different kits.

## Introduction

Interest in miRNAs has steadily increased since their discovery in the early 1990s due to their roles in diverse biological processes^1-4^ and their dysregulation associated with several diseases ^5-7^. Next generation sequencing (NGS) is an attractive technology to study miRNAs because of its high sensitivity and ability to detect novel miRNAs. Several commercially-available kits are available to prepare miRNA libraries for sequencing, which entails addition of adapter sequences to the miRNAs followed by reverse transcription and cDNA synthesis. In a recent study, we compared the performance of six such kits (CATS, CleanTag, QIAseq, TailorMix, SMARTer-beta and srLp) with respect to detection rate sensitivity, reliability and ability to detect differentially expressed miRNAs ^8^. However, two commonly used kits (NEBNext and NEXTflex) were not included.

Previous studies have reported differences in miRNA abundance detected by sequencing relative to the original RNA sample, which makes miRNA quantification challenging ^9 10^. Sequencing library preparation, and in particular the adapter ligation steps, have been identified as the primary sources of this bias ^10 11^. Most kits utilize RNA ligases to attach adapters to the miRNAs (e.g. NEBNext, QIAseq, TailorMix, CleanTag) but the efficiency of this step depends on the ligase used, the adapter sequence and the primary and secondary structure of the miRNA ^10-13^. NEXTflex reagents attempt to increase efficiency and reduce bias at this step by utilising adapters containing stretches of random nucleotides, which increases adapter sequence diversity. Other attempts to avoid bias whilst introducing adapter sequences onto miRNAs are polyadenylation and template switching oligonucleotides (e.g. CATS) or by using single adapter circularization (e.g. SMARTer).

In this study, we investigated the performance of the NEBNext and NEXTflex kits (Table 1). It should be noted that although the study aimed to test low input kits handling inputs below 100 ng, the NEBNext kit is not designed for inputs below 100 ng, but was nonetheless included as it is widely used. Both studies were performed under the same conditions with one exception: While in the first study the library preparation was performed by the kit vendors themselves, for the two kits presented in the present study this step was performed at Oslo University Hospital. This manuscript gives an overview on the results for all eight kits, with a focus on the NEBNext and NEXTflex kits. For more details on the other six kits we refer to Heinicke, et al. ^8^.

**Table 1:** Small RNA library preparation methods tested in this study.

| Kit name | Commercial supplier | Key features* | Max. input volume tolerated | Reported RNA input range (varies with type of input tested) | Max. number of indexes available | Method types |
| --- | --- | --- | --- | --- | --- | --- |
| NEXTFLEX® Small RNA-Seq Kit v3 (NEXTflex) | PerkinElmer Inc. | 5-step process of 3' adapter ligation, adapter inactivation, 5' adapter ligation, reverse-transcription and PCR. 3 purification steps. | 10.5 µl | Total RNA (1 ng - 2 µg), purified small RNA (from 1 - 10 µg total RNA), and a synthetic miRNA pool (≥100 pg) | 96 | Ligase based. 2-adapter procedure. Utilizes adapters with randomized 4mer ends |
| NEBNext® Small RNA Library Prep (E7300) (NEBNext) | New England Biolabs Inc. | Single-tube, 5-step process of 3' adapter ligation, primer annealing, 5' adapter ligation, reverse-transcription and PCR. 1-2 purification steps. | 6 µl | Total RNA (100 ng-1 µg) | 48 | Ligase based. 2-adapter procedure |

\* A step is defined as a labwork period that culminates in an incubation longer than 5 minutes.

## Results

Altogether 21 samples, comprising 15 synthetic miRNA samples (five mixes processed in triplicates) and six human total RNA samples (pool of rheumatoid arthritis patients and pool healthy controls processed in triplicates), were used to assess the performance of the different library preparation kits (Figure 1A). To aid comparison we present here the results of all eight kits, with our previous results ^8^ displayed in faded colours in the figures. Following library preparation, the NEBNext and NEXTflex libraries were sequenced together (i.e. on the same sequencing flow cell) with the libraries from the other six library preparation kits^8^. For NEBNext and NEXTflex, cluster density and read numbers passing filters were similar to the other kits that previously performed well (CleanTag, QIAseq, srLp, TailorMix) (Supplementary Figure 1 and Table 2).

**Figure 1:**
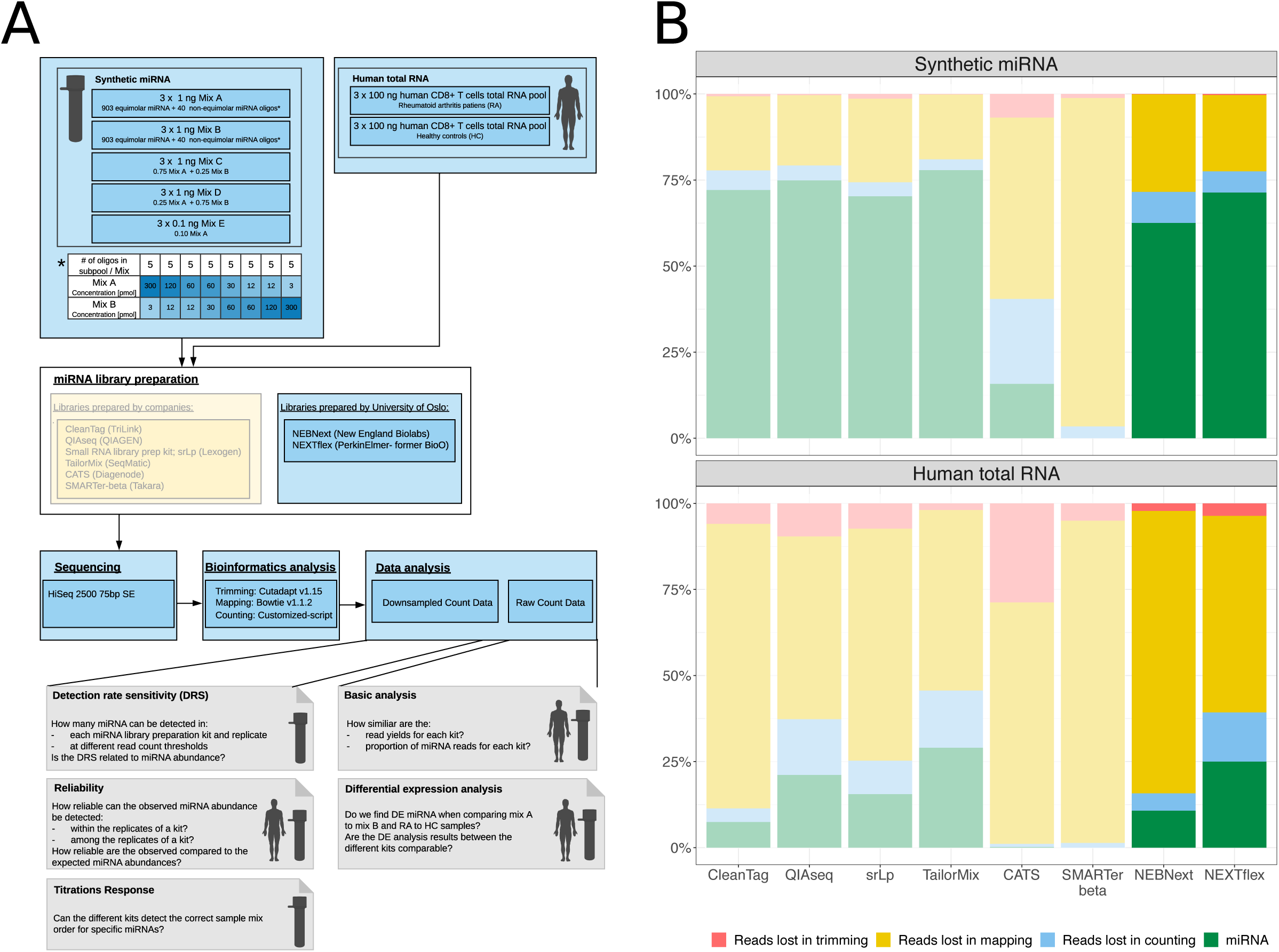
Experimental design and sequencing read distribution. (A): Overview of the study material, miRNA library preparation kits used, sequencing, bioinformatics and data analysis. Library preparation was performed in house in contrast to the study design presented in ^8^. Grey boxes represent individual data analysis steps. (B): Percentage of reads that were removed during the bioinformatic analysis and final miRNA proportion remaining (green). Trimming refers to removal of adapter sequences, mapping to miRNA reference alignment, and counting to filtering of aligned miRNAs that did not have the same length as the reference sequence. Results presented are the mean of 15 replicates in the synthetic miRNA (upper panel) and the mean of six replicates in the human total RNA samples (lower panel). Faded colors were used to indicate previous results ^8^. Images from Servier Medical Art (Servier. www.servier.com, licensed under a Creative Commons Attribution 3.0 Unported License) were used in (A).

Consistent with our earlier study, the greatest proportion of reads, both for NEBNext and NEXTflex, were discarded during mapping to the miRNA reference sequences (Figure 1B, Supplementary Figure 2 and Supplementary Table 1). Notably, the NEXTflex kit compared favourably to the best performing kits identified previously, and despite not being designed to handle sub-100 nanogram amounts, NEBNext performed adequately. To comprehensively evaluate the performance of the kits, read numbers were randomly down-sampled (2.5 million reads for synthetic miRNA samples and 0.75 million for human total RNA samples) and, where stated, were regularized log (rlog) transformed for subsequent analysis steps.

To assess the detection rate sensitivity of the library preparation kits, we tested several detection thresholds in the down-sampled synthetic miRNA samples. First, we defined a miRNA to be detected in a sample when at least one read in toal was registered. NEXTflex detected 928-934 of 943 miRNAs across all three replicates of the different synthetic mixes while NEBNext detected between 869-881 miRNAs (Figure 2A). Compared to our previous results, NEBNext was the kit that detected fewest miRNAs in all replicates of the different mixes. Furthermore, in mix E, where the RNA input was 10 times lower than in mixes A to D, NEBNext detected the fewest number of miRNAs across all three replicates. In contrast, NEXTflex, together with QIAseq and TailorMix missed the fewest miRNAs in one, two or all three replicates. The undetected miRNAs were generally kit specific (Supplementary Figure 3). However, some miRNAs such as EBV-1-3P and MIR-612, EBV-20-3P, MIR-548D-3P and MIR-193A-3P (miRNA annotation according to miRXplore Reference) were undetected across several kits and replicates (Supplementary Figure 4).

**Figure 2:**
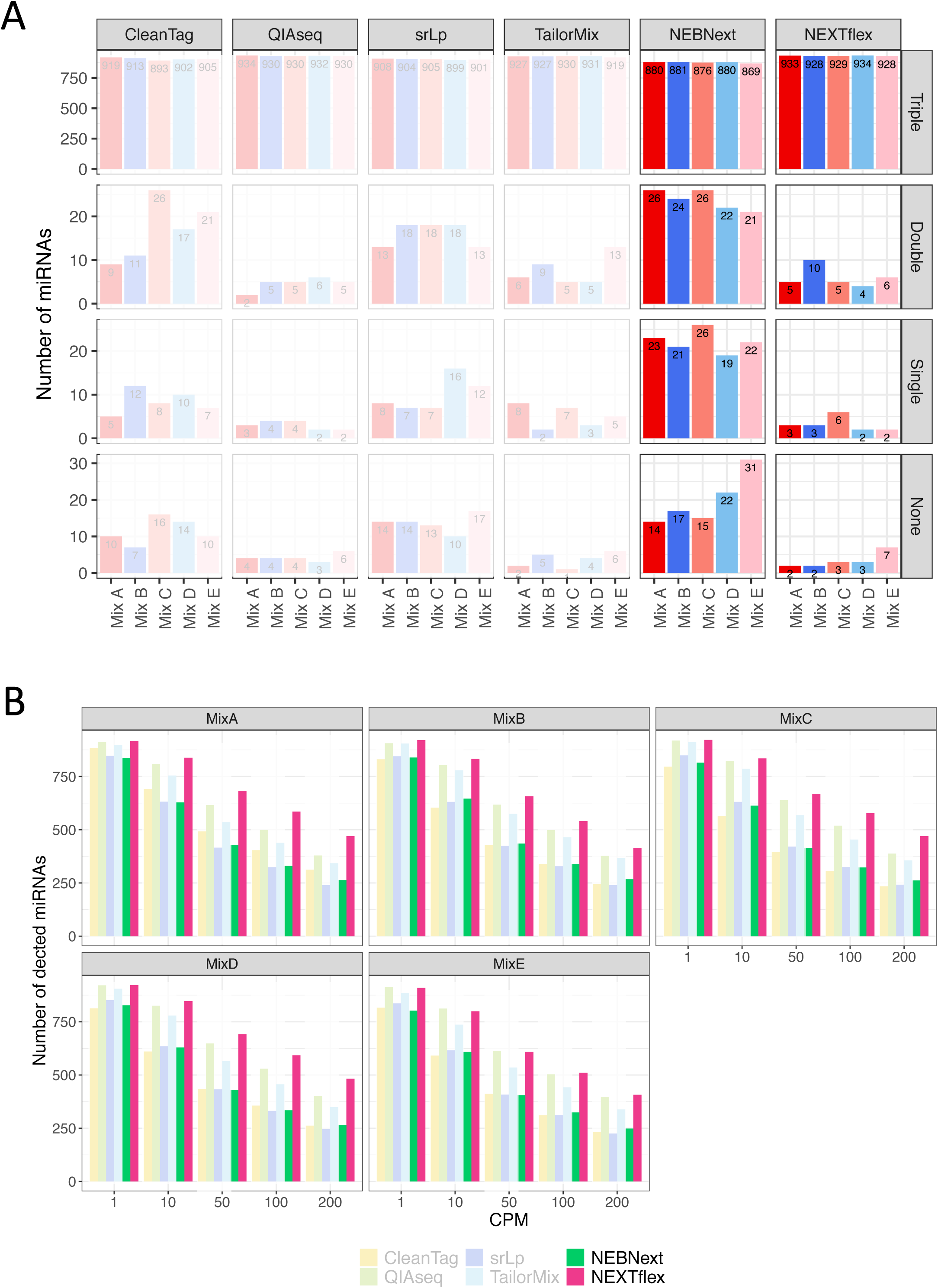
Detection rate sensitivity. (A): Bar charts presenting number of miRNAs detected in all replicates (Triple), in 2 out of 3 replicates (Double), in 1 out of 3 replicates (Single) or not detected in any replicate (None) across all synthetic miRNA mixes and all library preparation kits. The maximum number of detectable miRNAs is 943 (903 equimolar and 40 non-equimolar miRNA). (B): Bar charts for various read count thresholds in the synthetic miRNA samples. A miRNA is defined as detected when it is (i) expressed in all three replicates of the mix and (ii) the read counts are greater or equal to the count per million (CPM) threshold displayed on the x-axis. Faded colors were used to indicate previous results ^8^.

When analysing the 40 non-equimolar miRNAs, NEXTflex revealed a very high detection rate sensitivity, second only to the previously tested QIAseq kit (Supplementary Figure 5). Conversely, for NEBNext we observed the lowest detection rate sensitivity (except for the CATS and SMARTer-beta kits which were excluded from the analysis at this step already). Most of the miRNAs that could not be detected were present at low concentration levels. However, miRNA detection was not solely dependent on the concentration level (Supplementary Figure 5), suggesting that kit-specific biases also play a role.

Next, we examined sensitivity under more stringent detection thresholds, requiring a miRNA to be detected when at least 1, 10, 50, 100 or 200 read counts per million (CPM) were registered across all three mix replicates. With the exception of the non-equimolar miRNAs presented at the lowest concentration levels, all synthetic miRNAs should theoretically be detected at 200 CPM. However, as observed previously in Heinicke, et al. ^8^, the number of detected miRNAs decreased greatly with increasing CPM threshold for the NEXTflex and NEBNext kits (Figure 2B). For mix A to D, NEXTflex detected the most miRNAs among all tested kits while the detection sensitivity was similar to the QIAseq kit for mix E. NEBNext detected fewer miRNAs across all mixes and CPM thresholds than NEXTflex and obtained similar results to CleanTag and srLp.

We used down-sampled and rlog transformed miRNA count data to assess reliability. The intra-rater reliability (miRNA read count concordance within the replicates of a library preparation kit) of NEBNext and NEXTflex were as strong as for the previously tested kits, although slightly weaker results were observed for mix E with NEBNext. Both kits revealed ICC values between 0.93 and 0.99 (Supplementary Table 2) and Pearson correlation coefficients above 0.91 (p < 0.05, Supplementary Table 3). Bland-Altman plots (data not shown) indicated no systematic differences in the measurements.

To examine inter-rater reliability (miRNA read count concordance between the library preparation kits) the first replicate of each synthetic miRNA mix, rheumatoid arthritis (RA) or healthy control sample from all six library preparation kits (NEBNext, NEXTflex, CleanTag, QIAseq, TailorMix, srLp) was chosen. Larger differences were observed between the different library preparation kits than within the replicates of a kit with regard to miRNA reads counts. Similar to our previous study, ICC values were above 0.8 for the synthetic miRNA sample mixes and above 0.95 for the RA or healthy control samples (Supplementary Table 4). The same was true for the Pearson correlation coefficients which were above 0.73 and 0.92 (p < 0.05) for the synthetic miRNA and human total RNA samples respectively (Supplementary Table 5). No systematic differences in the measurements were observed by Bland-Altman analysis (data not shown).

As a further assessment of reliability, we investigated the concordance between the theoretical miRNA concentrations and the obtained read counts for the synthetic miRNA samples. For the 903 equimolar miRNAs, no significant deviation between a specific miRNA rlog read count and the median rlog read count over all equimolar miRNA was expected to be seen. The fold deviation was defined to be equimolar when its absolute value was less or equal to one. However, for the randomly chosen first replicate of mix A, only between 37.2% to 42.6% of the miRNAs were detected as equimolar. NEBNext detected the lowest number miRNAs to be equimolar while NEXTflex detected the highest number across all tested kits (Supplementary Figure 6).

To compare the performance of the kits for quantifying miRNA levels, the read counts of the 40 non-equimolar miRNAs were correlated with the expected theoretical levels. NEXTflex showed slightly lower correlations across all samples than QIAseq, which obtained the highest correlation coefficients in our previous study (Supplementary Table 6 and Supplementary Table 8 in ^8^). NEBNext was a middle-ranking kit in this correlation. However, as before, we found that none of the tested kits could accurately quantify the majority of miRNAs.

To examine kit performance in differential miRNA expression, non-down-scaled and untransformed miRNA counts were analyzed. Between mix A and mix B of the synthetic miRNA samples, 29 out of 40 differentially expressed miRNAs were detected by NEBNext and 26 by NEXTflex (Figure 3A). In comparison, all previously tested library preparation kits were able to detect between 32 to 35 differential expressed miRNAs. However, of those not all miRNAs were true positives. While only differentially expressed miRNAs were expected to be found within the pool of non-equimolar miRNA (n=40), an additional one to two equimolar miRNAs were detected to be differentially expressed by the previously tested library preparation kits. This was not the case for NEBNext or NEXTflex. MiRNAs that could not be detected as differentially expressed between mix A and B were often those with the lowest concentration level differences (Figure 3C). Quantitative reverse-transcriptase PCR assays on 16 of the 40 non-equimolar miRNAs revealed that the intended ratios for mix A and mix B were as expected (Supplementary Figure 7).

**Figure 3:**
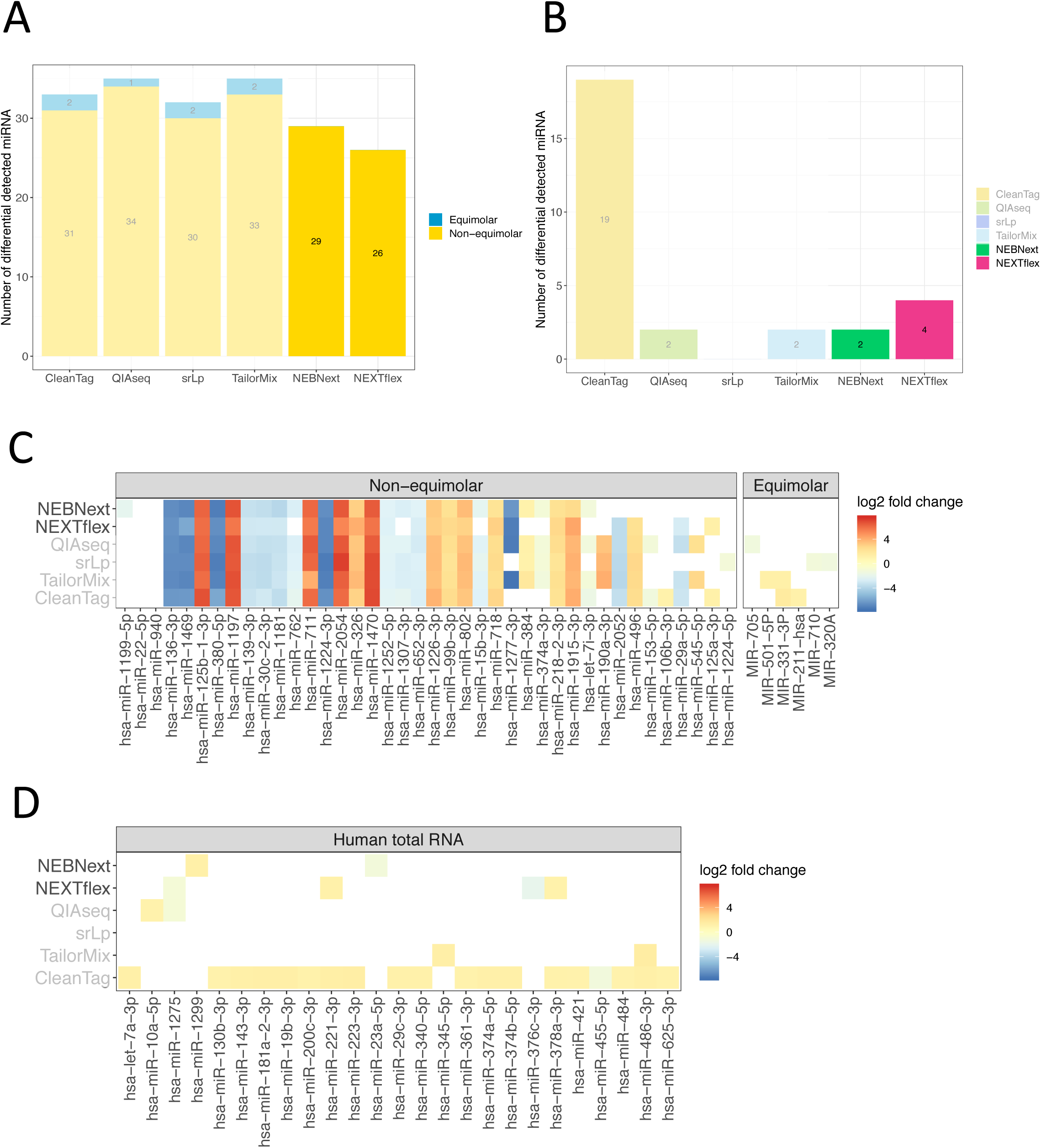
Differential expression analysis. Kit-specific number of differentially expressed miRNA detected for (A): synthetic miRNA samples (mix A versus mix B) and (B): human total RNA samples (RA versus healthy control). miRNA-specific log2 fold changes across the different kits for (C): synthetic miRNA samples and (D): human total RNA samples. Faded colors or grey font were used to indicate previous results ^8^.

We also performed differential expression analysis between the RA patient and healthy control pools of human total RNA samples. NEBNext detected two and NEXTflex four significant differentially expressed miRNAs (Figure 3B), but the kits did not identify the same miRNAs as differentially expressed. There was also no overlap between the differentially expressed miRNAs predicted by NEBNext and those predicted by the previously-tested miRNA library preparation kits. For NEXTflex, three of the four miRNAs were already previously detected as differentially expressed by other kits ^8^: hsa-miR-1275 was also detected by QIAseq to be down-regulated in RA patients compared to healthy controls while hsa-miR-378a-3p and hsa-miR-221-3p were detected by CleanTag to be up-regulated in RA patients versus healthy controls (Figure 3D).

Finally, we compared the performance of the kits in the titration response assay, which provides a measure of quantitative performance ^14 15^. Downscaled and rlog transformed read counts of the 40 non-equimolar miRNAs were scored for their adherence to expected concentration orders in mixes A-D, with five miRNAs in each of the eight concentration groups (Table 3). In this assay, NEBNext performed better than NEXTflex, which had an intermediate performance relative to the results reported previously.

**Table 2:** Median and standard deviation (SD) of the raw read counts passing sequencing quality filters for each kit and sample type.

| Kit | Sample Type | Median | SD |
| --- | --- | --- | --- |
| NEBNext | synthetic miRNA | 15963032 | 2098564 |
|  | human total RNA | 12945516 | 934152 |
| NEXTflex | synthetic miRNA | 15726206 | 3428519 |
|  | human total RNA | 9947511 | 865005 |
| CATS | synthetic miRNA | 1657065 | 1686647 |
|  | human total RNA | 4368917 | 610984 |
| srLp | synthetic miRNA | 21708163 | 3074872 |
|  | human total RNA | 9553164 | 3234006 |
| QIAseq | synthetic miRNA | 25025406 | 4866588 |
|  | human total RNA | 17161083 | 1492933 |
| TailorMix | synthetic miRNA | 12904412 | 2208956 |
|  | human total RNA | 11875567 | 1275394 |
| SMARTer | synthetic miRNA | 4817693 | 2249898 |
|  | human total RNA | 714966 | 296656 |
| CleanTag | synthetic miRNA | 10044117 | 2055836 |
|  | human total RNA | 19647913 | 4898198 |

**Table 3:** Fraction of titrating miRNAs (n = 5) in each of the eight concentration groups. Average rlog expression values for the 40 non-equimolar miRNAs were calculated across the three replicates each of mix A to D. Each miRNA was scored as titrating if the average values followed the expected trend in concentrations from high to low or vice versa across mixes A to D. Grey font indicates previous results ^8^.

| Conc.<br>Ratio | NEBNext | NEXTflex | CleanTag | QIAseq | srLp | TailorMix |
| --- | --- | --- | --- | --- | --- | --- |
| 0.01 | 1 | 0.8 | 1 | 1 | 1 | 1 |
| 0.1 | 1 | 0.6 | 0.8 | 1 | 1 | 1 |
| 0.2 | 0.8 | 1 | 1 | 1 | 0.8 | 0.8 |
| 0.5 | 0.8 | 0.6 | 0.8 | 0.6 | 0.4 | 0.6 |
| 2 | 0.8 | 0.6 | 0.6 | 0.8 | 0.8 | 0.2 |
| 5 | 1 | 1.0 | 0.4 | 1 | 1 | 0.8 |
| 10 | 0.6 | 0.8 | 0.6 | 1 | 1 | 0.6 |
| 100 | 0.8 | 0.8 | 0.8 | 1 | 0.8 | 0.8 |

## Discussion

We assessed the performance of NEBNext and NEXTflex and present the results along with the six library preparation kits we tested previously ^8^. Identical RNA input samples prepared at the same time point and under the same conditions were used in both studies. The prepped sample libraries from all kits were sequenced on the same flow cell and identical bioinformatics and data analysis steps were performed. However, the studies differ in the way in which the library preparation was performed: While it was performed by the kit vendors themselves in our first study^8^, we performed library preparation for this additional study. Although our aim was to make the two studies as similar as possible, we cannot exclude that the different library preparation approaches may have influenced the results. In the present study, researchers experienced with library preparation performed the work, therefore, the outcome for NEBNext and NEXTflex may represent results that can be obtained by an “average” user. In contrast, in our previous study where the library preparation was performed by the vendors, it was expected the results represent best-case-scenarios. Furthermore, since the datasets for NEBNext and NEXTflex were generated from individual sequencing lanes, unlike for most kits in the first study which were distributed across several lanes, we cannot exclude that lane-specific effects on data quality may have influenced the conclusions in this current work.

Jayaprakash, et al. ^11^ showed that small RNA profiles are dependent on the adapter sequences used during library preparation and according to their recommendation a mix of adapters will enable more accurate estimation of miRNA abundance. NEXTflex is the only tested kit in our study that uses this approach by including randomized adapter termini in the procedure. Compared to the three fixed-adapter kits (NEBNext, srLp and CleanTag), the overall performance of NEXTflex with respect to detection rate sensitivity, reliability and differential expression was superior. However, QIAseq and TailorMix also used fixed adapters and performed slightly better than or equally as well as NEXTflex. Even though including randomized adapter sequences during library preparation seems to improve the performance of a kit, our study suggests that additional factors influence the performance. These factors might include, for example, type of ligase or ligation temperature and ligation time. Giraldez, et al. ^16^ have also suggested that the concentration of polyethylene glycol during the ligation reactions affects performance, but since buffer constituents provided by commercial vendors are kept proprietary, we were unable to examine this parameter.

With the exception of the titration response assay, NEXTflex generally displayed one of the best performances, whilst NEBNext showed average performance. In particular, the NEBNext kit displayed lower miRNA detection sensitivity than the other kits. This was especially evident for the synthetic miRNA mix E. In this mix NEBNext detected the lowest number of miRNAs across all kits and mixes. The analysis of the non-equimolar miRNAs revealed that miRNAs with low abundance often remained undetected by NEBNext, and its reliability was lower on mix E. According to the NEBNext manual, the kit allows a minimum input of 100ng total RNA. MixE had the lowest miRNA content (0.1 ng in 10 ng total RNA) thus it is not surprising that NEBNext showed poorer detection sensitivity compared to the other library preparation kits. However, some of the miRNAs remained undetected independent of their abundance levels which indicates that additional factors influence their detection and therefore the kit performance. This is true for all tested kits: i.e. the kits appear to have preferences for certain miRNAs. It was previously suggested that the terminal nucleotides of the miRNAs influence their detection^9^ as well as the secondary structure of the miRNA ^17^ and co-folding between miRNA and adapter^12^, which may explain the kit-specific preferences observed.

Both the NEXTflex and NEBNext kits detected fewer differentially expressed miRNAs than the kits reported previously. Whilst this is not surprising for the NEBNext kit, which appears to be less sensitive, it was unexpected or the NEXTflex kit. However, this lower sensitivity was balanced by fewer false positive calls, which might be of advantage for studies interested in finding novel biomarkers for e.g. specific diseases or treatment responses where false positives are particularly undesirable.

In conclusion, we found considerable differences between the library preparation kits when comparing their performance. Overall, QIAseq demonstrated the best performance followed by TailorMix and NEXTflex. NEBNext, srLp and CleanTag were ranked as medium performance kits. However, when it comes to accurate quantification of miRNA, all tested kits show room for improvement.

## Material and Methods

The study material was described in detail in Heinicke, et al. ^8^. Briefly, synthetic miRNA and human total RNA samples were used as input into library preparation. The performances of a total of eight kits (six kits from our previous and two kits from the present publication) were compared using triplicate samples as summarised below and in Figure 1A). Synthetic miRNA samples consisted of equimolar (n=962, miRXplore Universal Reference, Miltenyi, California, United States) and non-equimolar miRNA oligonucleotides (n=40, Eurofins MWG Synthesis GmbH, Bavaria, Germany) which were used to create five different mixes, A-E. Mix A and B contained the same equimolar pool of miRNAs, but differed in eight concentration ratios of the 40 non-equimolar miRNAs (Supplementary Table 1 in ^8^). Mix C was a 0.75 titration of mix A and 0.25 titration of mix B while the titration ratio for mix D was vice versa. Mix E equates mix A but at a 10-fold lower concentration. *Saccharomyces cerevisiae* total RNA was added to the different mixes to obtain a more complex RNA mixture. In each mix the RNA content was 2 ng/ul and miRNAs represented approximately 10% (w/w) in mix A to D and 1% (w/w) in mix E (Supplementary Table 2 in ^8^). The intended mix ratios were verified using RT-qPCR with 16 pre-designed TaqMan Small RNA assays (Thermo Fisher Scientific, Massachusetts, United States, Supplementary Material and Methods in ^8^).

Human total RNA samples were extracted from peripheral blood CD8+ T cells from a pool of either newly diagnosed rheumatoid arthritis (RA) patients (n=4) or healthy controls (n=4). For all samples the RNA integrity value was above 8.5.

Library preparation for all kits except NEBNext and NEXTflex was described previously (see Supplementary Material and Methods section and Supplementary Table 2 in ^8^). NEBNext and NEXTflex libraries were prepared from the 21 samples described above according to manufacturer’s instructions. For the synthetic miRNA mix A to D, containing 10 ng miRNA oligonucleotides, NEBNext adapters were not diluted while NEXTflex adapters were diluted 1:2. For the synthetic miRNA mix E, containing 1 ng miRNA oligonucleotides, and the human total RNA samples the adapters were diluted 1:2 for NEBNext and 1:4 for NEXTflex. Synthetic miRNA samples mix A to D were amplified using 12 PCR cycles for NEBNext and 16 PCR cycles for NEXTflex while synthetic miRNA samples mix E and human total RNA samples were amplified using 15 PCR cycles for NEBNext and 20 PCR cycles for NEXTflex. TapeStation 2200 High Sensitivity D1000 reagents (Agilent Technologies, California, USA) were used to verify the presence of miRNA library constructs at approximately 143 bp for NEBNext and 150 bp for NEXTflex. Pippin Prep (Sage Science, Massachusetts, USA) with 3% Agarose Gel Pippin Cassettes was used to removed adapters dimers and other unwanted fragments. Per lane of the Pippin Cassette five to six samples were pooled together. Size selection was optimized to cover fragments from ca 130bp to 160bp. Final library yields and size were measured on a Bioanalyzer 2100 using high sensitivity reagents (Agilent Technologies, Supplementary Figure 8).

Libraries were sequenced on one single-read flow cell of a HiSeq 2500 (Illumina, California, United States) with 75bp reads. Each of the 21 libraries from NEBNext and NEXTflex were sequenced independently from the previously tested library preparation kits on two single lanes (Supplementary Figure 9). Cutadapt ^18^ v1.15 was used to trim the following adapter sequences from the demultiplexed fastq files: AGATCGGAAGAGCACACGTCT (NEBNext) and TGGAATTCTCGGGTGCCAAGG (NEXTflex). For NEXTflex we additionally clipped the first and last 4 bases of the reads to remove the random 4mers that are included in the adapters. We found 59 oligonucleotide sequences from the miRXplore Reference to be identical to sequences in the yeast sacCer3 genome. Those sequences were removed from the synthetic miRNA reference to avoid downstream miRNA miscounting because of the yeast fragments (Supplementary Table 3 in ^8^). Trimmed reads were mapped without allowing for mismatches using bowtie ^19^ v.1.1.2 and counted using a customized script. The samples were randomly down-sampled to 2.5 million reads for the synthetic miRNA and 0.75 million reads for the human total RNA samples. To account for the heteroscedastic behaviour of miRNA-seq data, we transformed the count data using the rlog function of DeSeq2 ^20^ v1.20.0 where necessary.

Detection rate sensitivity was assessed by investigating which miRNAs could be detected in the synthetic miRNA samples using down-sampled read count data. The reliability of the different kits was investigated using rlog transformed downscaled data and assessing intra-rater correlation (ICC, two-way mixed model, absolute agreement and single rater), Pearson correlation and Bland-Altman agreements. Differential expression, using edgeR ^21^ v3.22.3, between mix A and B for the synthetic miRNA samples and RA patients and healthy controls was assessed using the original read count data. A miRNA was defined as significantly differentially expressed if the absolute value of the log fold change was above 1 after adjusting for multiple testing using the method of Benjamini and Hochberg, with a false discovery rate of 0.05. For the 40 non-equimolar miRNAs of the synthetic samples we assessed the titration response in mixes A-D using the average down-sampled rlog counts for each miRNA following the data analysis previously presented by Shippy, et al. ^14^. A miRNA was scored as titrating if its average expression value followed the expected concentration trend. Further details of bioinformatic analysis are given in ^8^.

Sequencing fastq files and miRNA count tables have been deposited in the Gene Expression Omnibus database with accession number GSE141658.

## Acknowledgments

We thank Iris Langstein and Philipp Korber for providing *S. cerevisiae* RNA. Images from Servier Medical Art (Servier. www.servier.com, licensed under a Creative Commons Attribution 3.0 Unported License) were used in Figure 1A.

## Appendices

**Supplementary Figures** (document: Supplementary_figures_Heinicke_etal2020.docx)

**Supplementary Tables** (document: Supplementary_tables_Heinicke_etal2020.docx)

