## Supplementary_Figures for "An extension to: Systematic assessment of commercially available low-input miRNA library preparation kits"

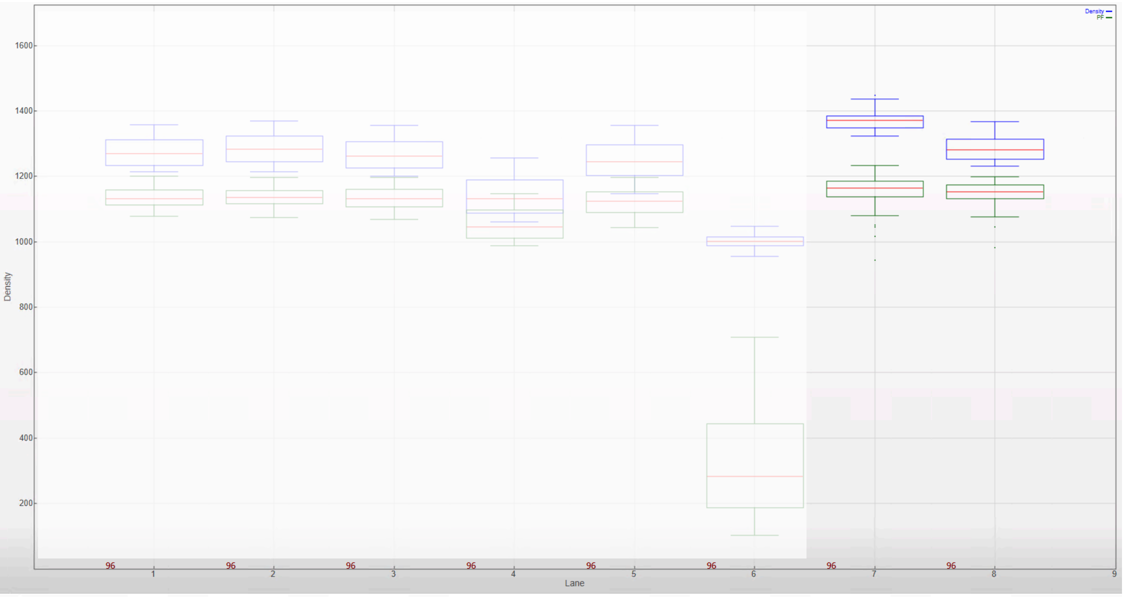


*Figure S1: Cluster density per lane of Illumina sequencing run. Image generated by Illumina Sequence Analysis Viewer software version 2.4.5, showing cluster density in thousands (K) of clusters per mm$^2$ of flowcell area for each sequencing lane. Raw cluster density is shown as blue boxplots, and clusters passing filters in green. Optimum raw cluster density as recommended by the manufacturer is 950-1050 K clusters per mm$^2$. Lanes 1-6 contain the libraries from the kits presented in our previous publication. Lane 7 contains libraries prepared with the NEXTflex and lane 8 contains libraries prepared with NEBNext.*

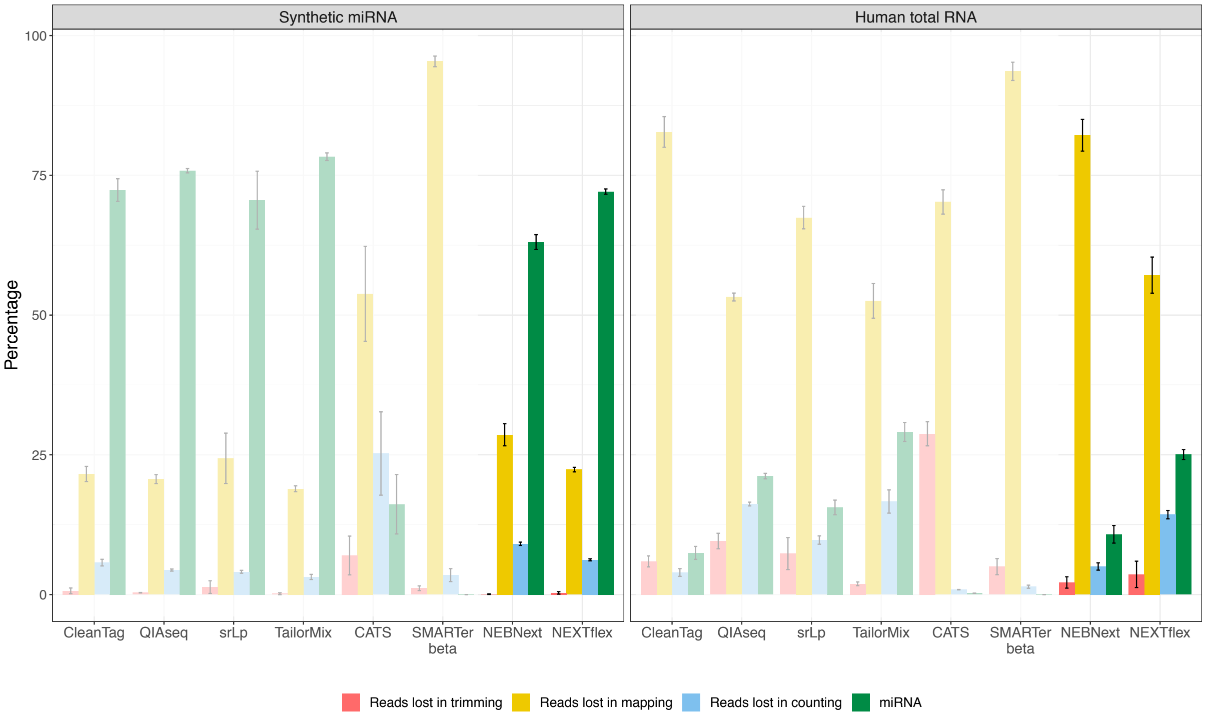


*Figure S2: Sequencing read distribution. Percentage of reads that were removed during the bioinformatics analysis and remaining miRNA reads. The results presented are the mean of 15 replicates in the synthetic miRNA and the mean of six replicates in the human total RNA samples. Error bars represent mean read count +/- standard deviation.*

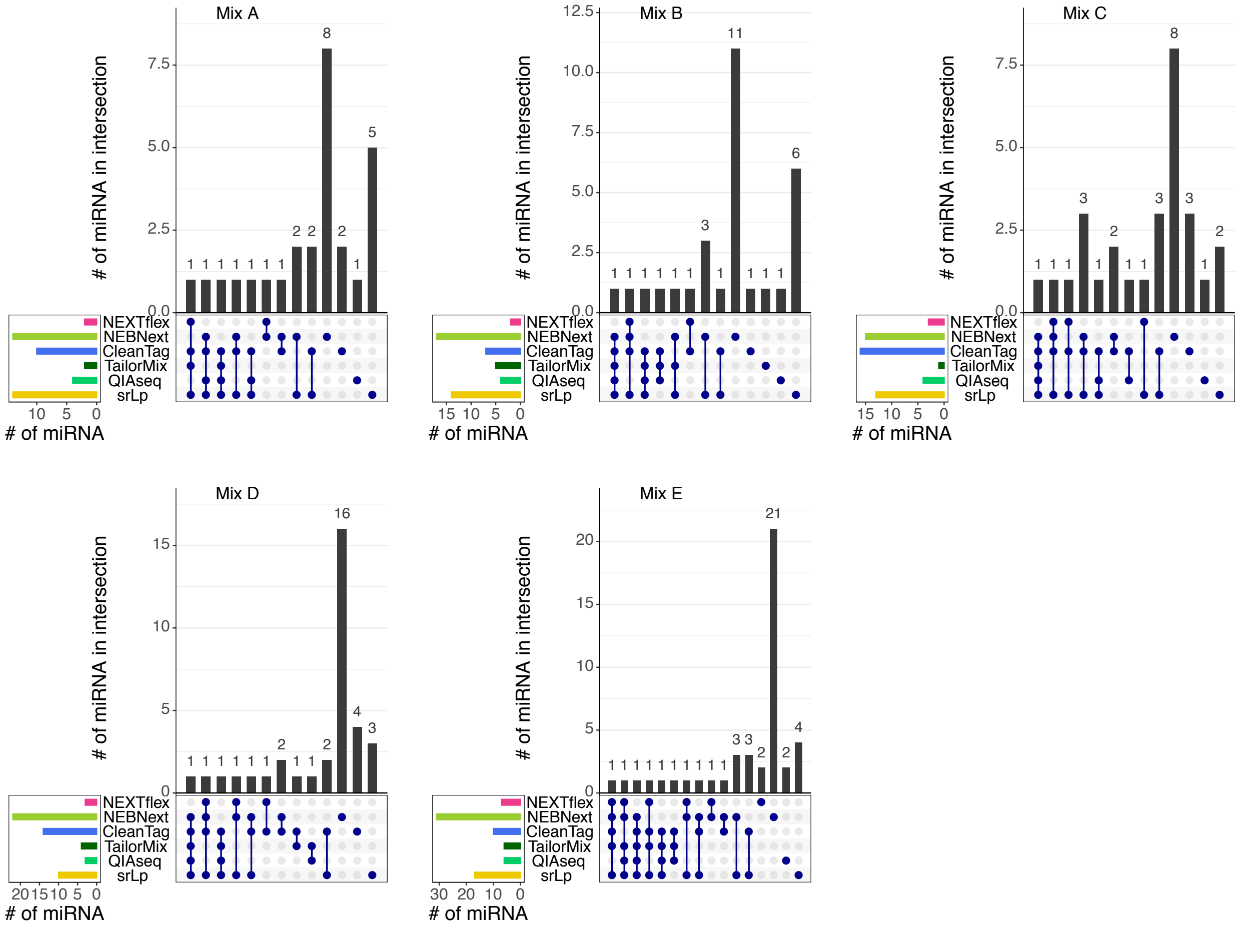


*Figure S3: Upset plots representing the intersections of the miRNA being undetected in all three replicates in a mix across the different kits and mixes. The interactions are indicated by the blue dots and lines. The number of miRNAs in the intersection is represented by the black bar charts.*

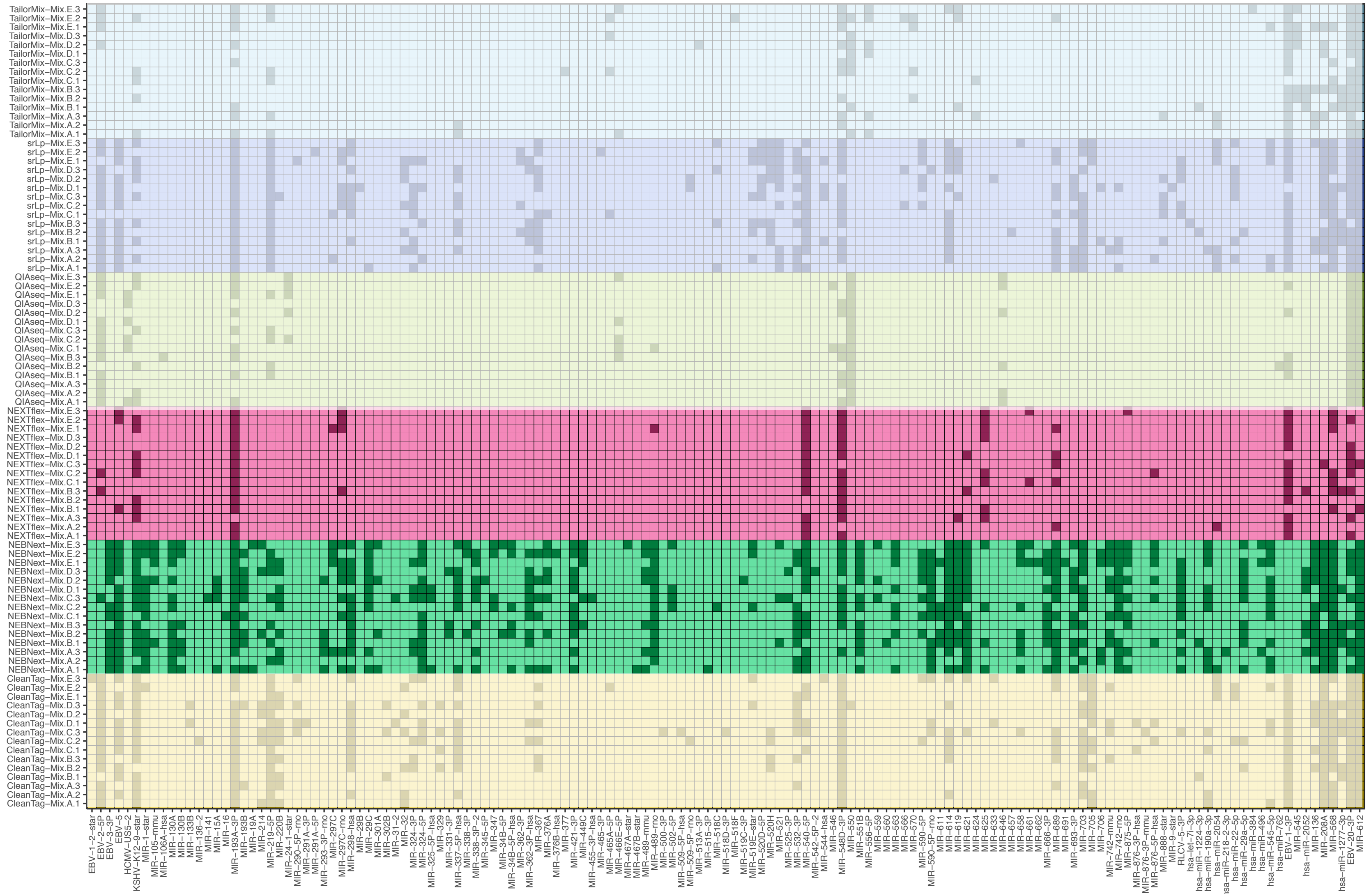

*Figure S4: Detection rate sensitivity for the synthetic miRNAs that could not be detected in at least one of the replicates for one of the kits and in at least one mix. The remaining miRNAs not represented in the plot were detected in all replicates of all kits. The library preparation kit and the replicate are presented on the y-axis. A darkened box within the plot indicates that the miRNA for this kit and replicate was not detected with at least one count.*

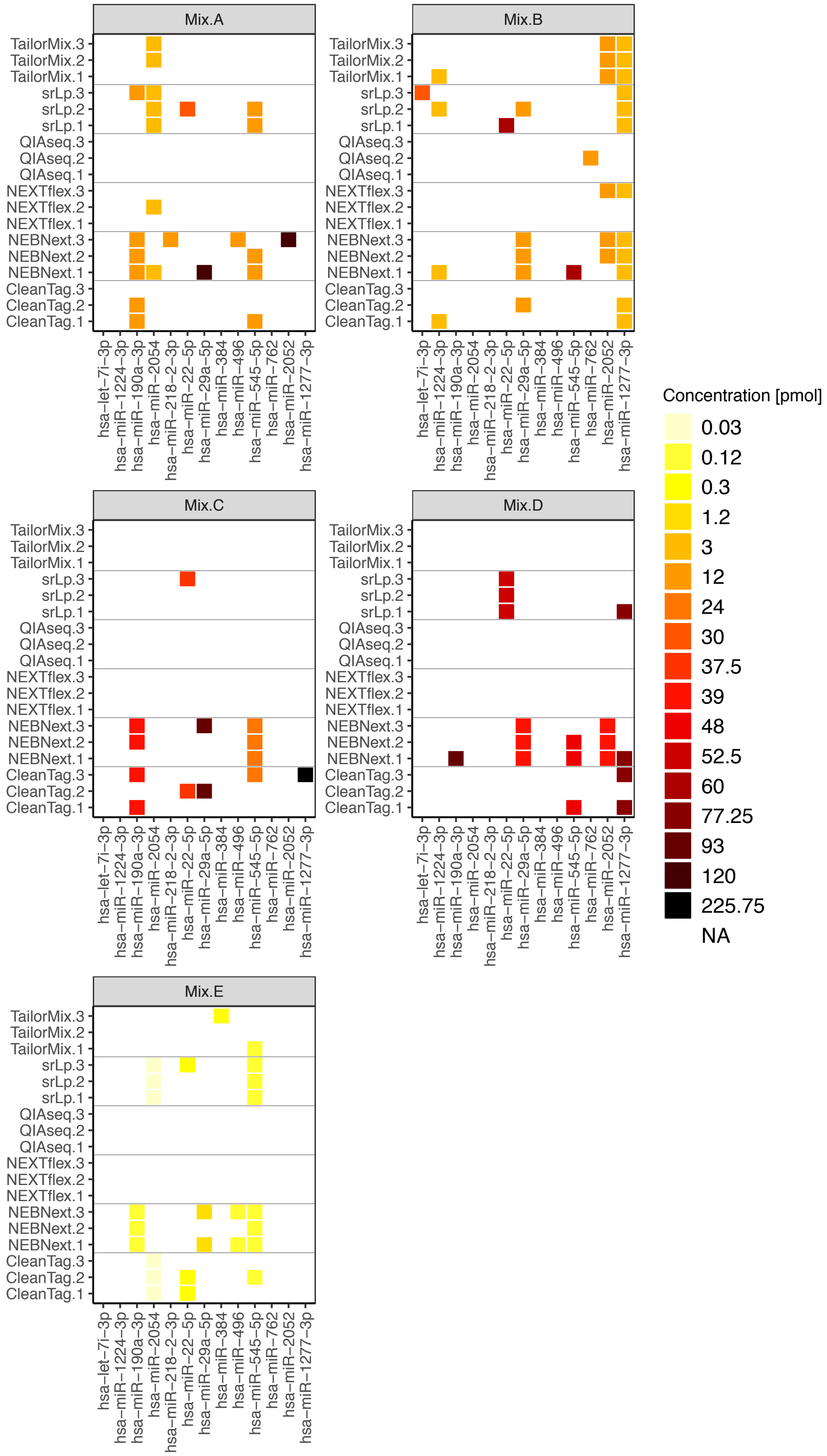


*Figure S5: Detection rate sensitivity. Each of the five subplots present the detection rate sensitivity for the non-equimolar miRNAs in mix A to mix E. miRNAs presented on the x axis (n=13) represent miRNAs that could not be detected in at least one of the replicates of one kit and in one mix. The remaining 27 miRNAs of the non-equimolar group (not presented) were detected in all replicates of all reagents in all five mixes. The library preparation reagents and the replicate are presented on the y-axis. A coloured box within a plot indicates that the miRNA for this kit and replicate could not be detected. The colour of the box represents the concentration of the miRNA.*

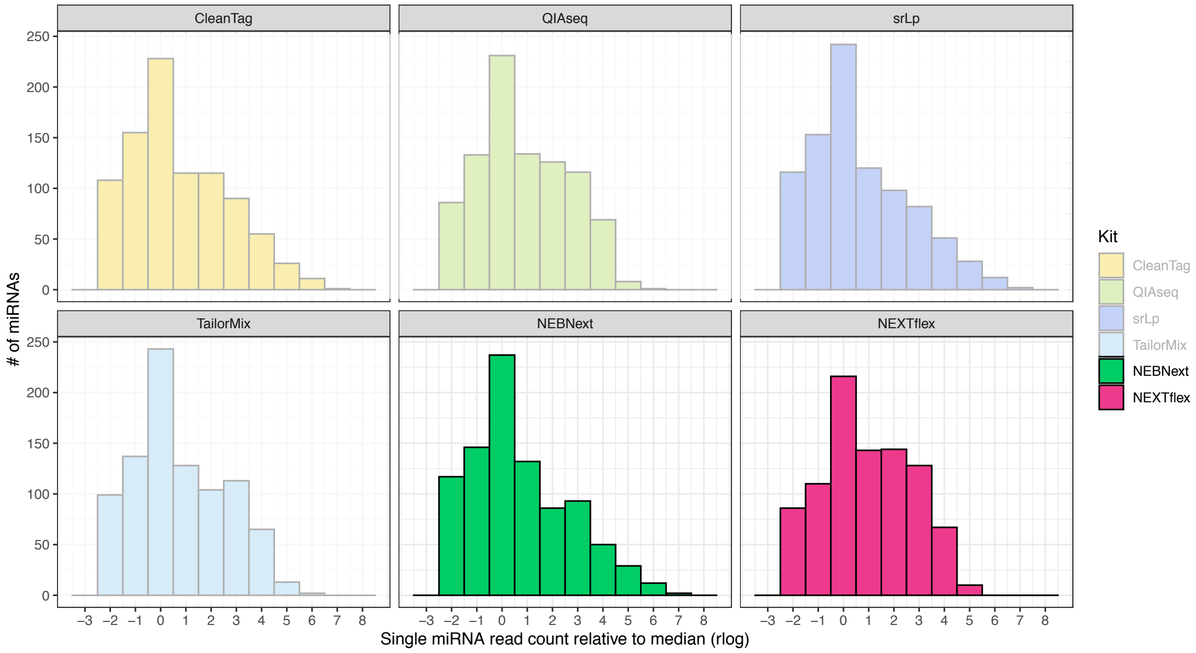


*Figure S6: Bias in miRNA detection utilizing the synthetic equimolar miRNA reference miRXplore. Bar charts represent the rlog ratio of single miRNA read counts to the median read counts of all the equimolar miRNAs of the specific library preparation kit. Results for replicate 1 of synthetic mix A are shown (near identical data was obtained for other replicates, data not shown).*

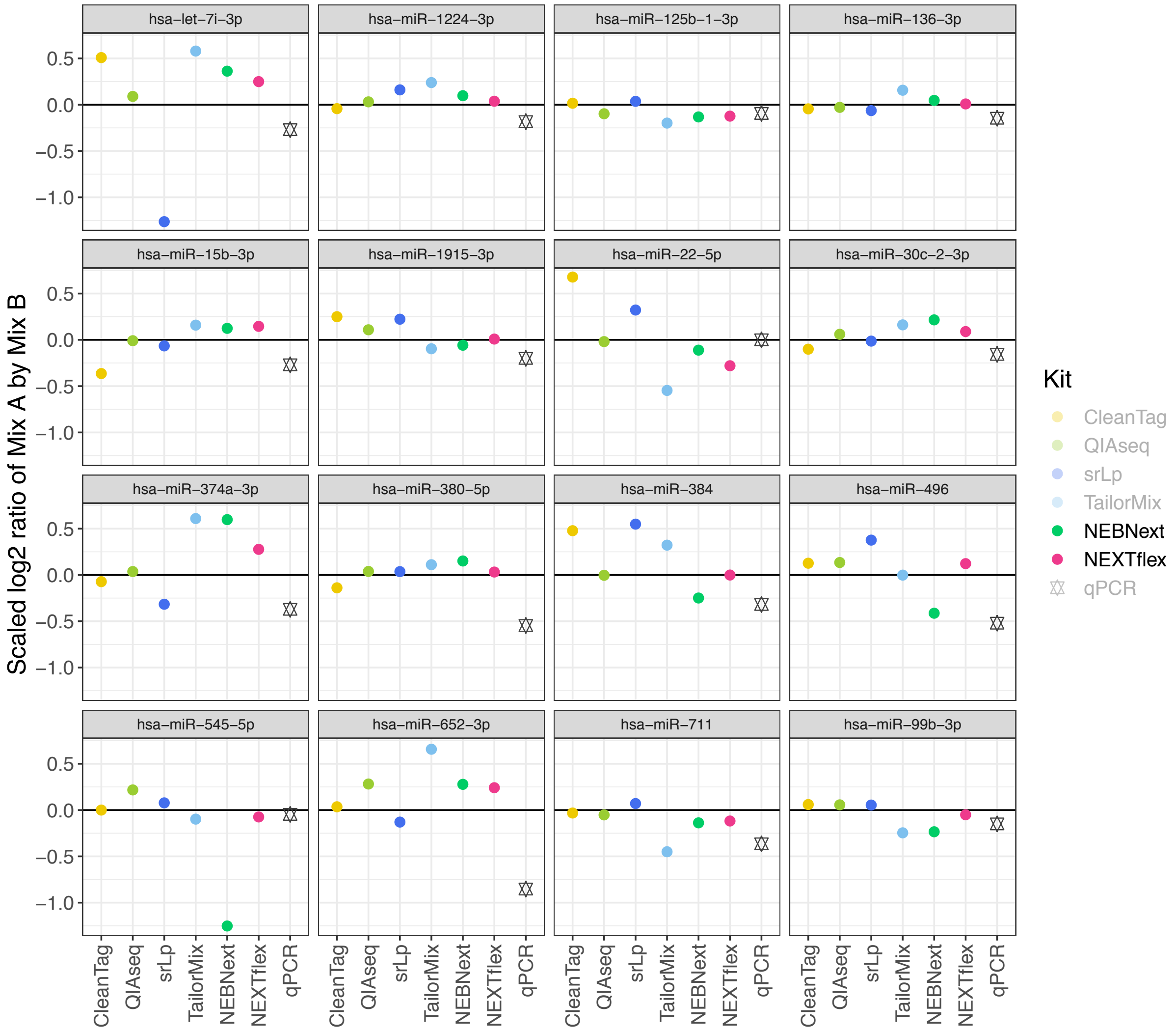


*Figure S7: Scaled and normalized log2 ratios of mix A and mix B for the read counts (sequencing data, represented by the library prep kits) and qPCR (copies detected relative to a standard curve) for 16 selected miRNAs. The horizontal black line represents the expected ratio.*

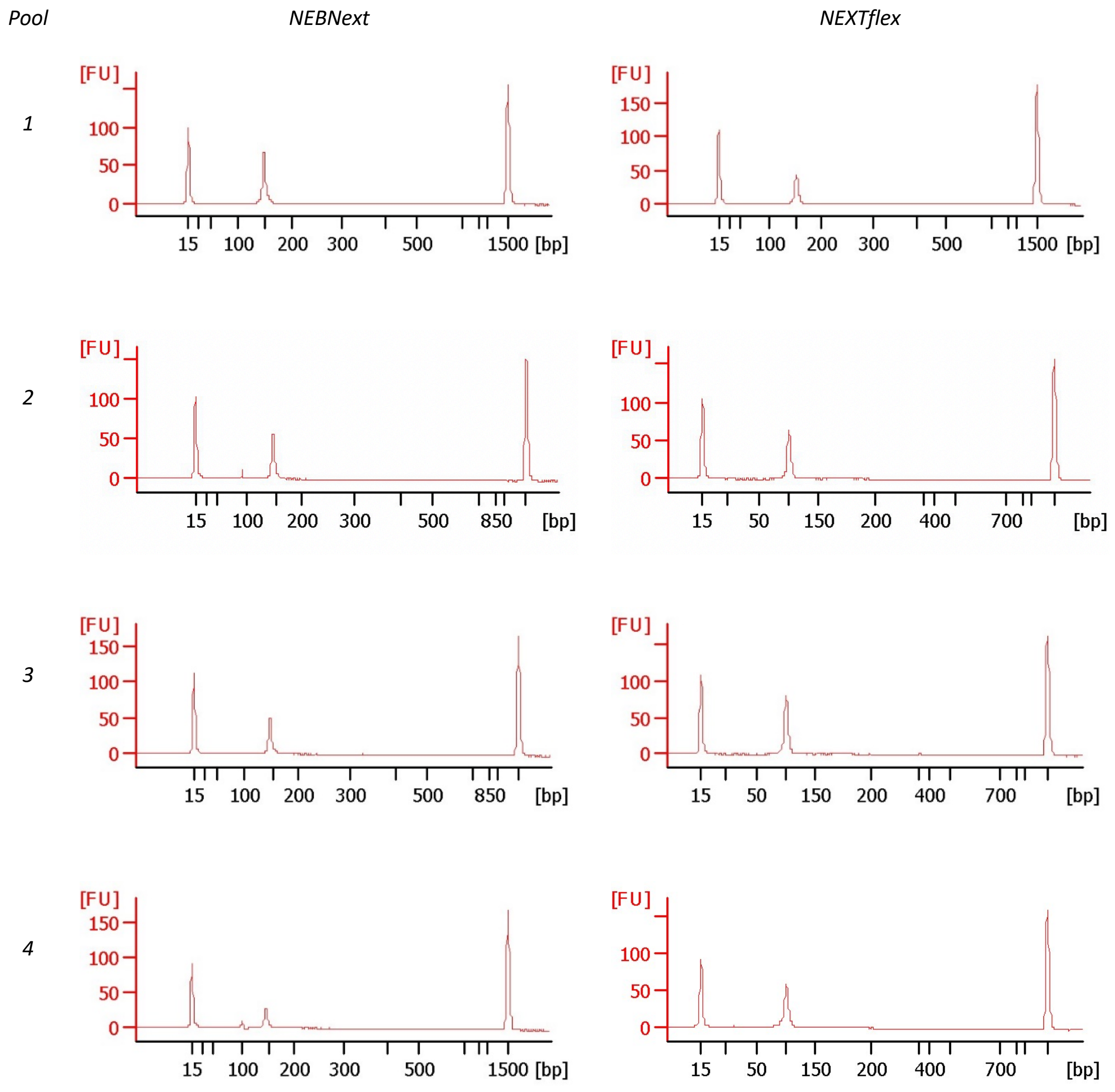


*Figure S8: Bioanalyzer electrophoretograms for pools of NEBNext and NEXTFlex libraries. X-axis in all cases is size in bp, and Y-axis fluorescence intensity (nucleic acid amount). Peaks at 15bp and 1500bp are internal reference size markers.*

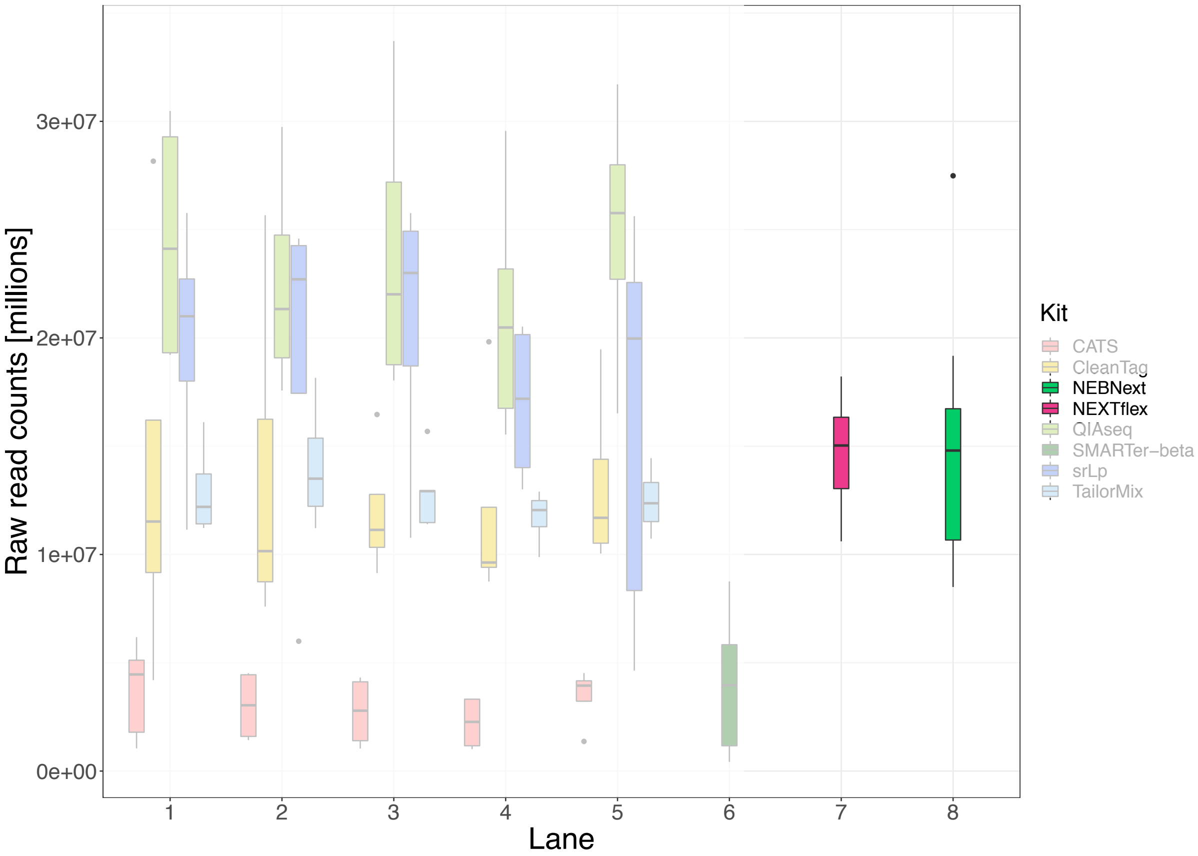


*Figure S9: Kit-specific raw read distributions obtained on eight lanes of a HiSeq2500 flow cell. Each lane consists of a pool of 21 libraries. Lanes 1-6 contain the libraries from the kits presented in our previous publication. NEXTflex libraries were sequenced separate from the other kits on lane 7 and NEBNext libraries on lane 8.*
