## Supplementary_Tables for "An extension to: Systematic assessment of commercially available low-input miRNA library preparation kits"

S1 Table

Overview of sample names, replicates, mix and the number of reads for the different bioinformatics steps

| Sample name | Kit_name | Kit_name_used_in_this_study | Mix | Replicate | HiSeq Lane | Raw_read_count | #_reads_after_trimming | #_reads_after_mapping | #_miRNA |
|---|---|---|---|---|---|---|---|---|---|
| BioO-1 | NEXTFLEX® Small RNA-Seq Kit v3 | NEXTflex | A | 1 | 7 | 18218249 | 18184936 | 14339500 | 13185753 |
| BioO-2 | NEXTFLEX® Small RNA-Seq Kit v3 | NEXTflex | A | 2 | 7 | 13991216 | 13969947 | 10974829 | 10099887 |
| BioO-3 | NEXTFLEX® Small RNA-Seq Kit v3 | NEXTflex | A | 3 | 7 | 12657727 | 12634626 | 9892733 | 9075767 |
| BioO-4 | NEXTFLEX® Small RNA-Seq Kit v3 | NEXTflex | B | 1 | 7 | 15963032 | 15938990 | 12529310 | 11544914 |
| BioO-5 | NEXTFLEX® Small RNA-Seq Kit v3 | NEXTflex | B | 2 | 7 | 15122884 | 15104390 | 11832735 | 10943067 |
| BioO-6 | NEXTFLEX® Small RNA-Seq Kit v3 | NEXTflex | B | 3 | 7 | 10605966 | 10562627 | 8201537 | 7557424 |
| BioO-7 | NEXTFLEX® Small RNA-Seq Kit v3 | NEXTflex | C | 1 | 7 | 16437557 | 16423076 | 12957132 | 11983328 |
| BioO-8 | NEXTFLEX® Small RNA-Seq Kit v3 | NEXTflex | C | 2 | 7 | 16336877 | 16311930 | 12916270 | 11898078 |
| BioO-9 | NEXTFLEX® Small RNA-Seq Kit v3 | NEXTflex | C | 3 | 7 | 16228788 | 16173940 | 12703971 | 11703634 |
| BioO-10 | NEXTFLEX® Small RNA-Seq Kit v3 | NEXTflex | D | 1 | 7 | 13071151 | 13034327 | 10135887 | 9345100 |
| BioO-11 | NEXTFLEX® Small RNA-Seq Kit v3 | NEXTflex | D | 2 | 7 | 15059341 | 15016546 | 11808742 | 10879255 |
| BioO-12 | NEXTFLEX® Small RNA-Seq Kit v3 | NEXTflex | D | 3 | 7 | 17180294 | 17152794 | 13494127 | 12431946 |
| BioO-13 | NEXTFLEX® Small RNA-Seq Kit v3 | NEXTflex | E | 1 | 7 | 14553886 | 14474756 | 11369332 | 10432151 |
| BioO-14 | NEXTFLEX® Small RNA-Seq Kit v3 | NEXTflex | E | 2 | 7 | 16780849 | 16631287 | 13091658 | 12027160 |
| BioO-15 | NEXTFLEX® Small RNA-Seq Kit v3 | NEXTflex | E | 3 | 7 | 18026411 | 17918629 | 14133034 | 12964861 |
| BioO-16 | NEXTFLEX® Small RNA-Seq Kit v3 | NEXTflex | RA | 1 | 7 | 12849397 | 12552787 | 5244011 | 3338687 |

| | | | | | | | | | |
|---|---|---|---|---|---|---|---|---|---|
| BioO-17 | NEXTFLEX® Small RNA-Seq Kit v3 | NEXTflex | RA | 2 | 7 | 12687019 | 12399919 | 4801880 | 3045581 |
| BioO-18 | NEXTFLEX® Small RNA-Seq Kit v3 | NEXTflex | RA | 3 | 7 | 15032359 | 14778245 | 5624252 | 3663029 |
| BioO-19 | NEXTFLEX® Small RNA-Seq Kit v3 | NEXTflex | HC | 1 | 7 | 13041634 | 12194118 | 5366906 | 3393604 |
| BioO-20 | NEXTFLEX® Small RNA-Seq Kit v3 | NEXTflex | HC | 2 | 7 | 12436643 | 11593045 | 4822958 | 3043765 |
| BioO-21 | NEXTFLEX® Small RNA-Seq Kit v3 | NEXTflex | HC | 3 | 7 | 13330601 | 13041586 | 5341258 | 3393106 |
| NEB-1 | NEBNext® Small RNA Library Prep | NEBNext | A | 1 | 8 | 27488251 | 27474709 | 19737504 | 17105707 |
| NEB-2 | NEBNext® Small RNA Library Prep | NEBNext | A | 2 | 8 | 14202691 | 14180351 | 10353602 | 8993620 |
| NEB-3 | NEBNext® Small RNA Library Prep | NEBNext | A | 3 | 8 | 13154785 | 13147059 | 9585597 | 8363552 |
| NEB-4 | NEBNext® Small RNA Library Prep | NEBNext | B | 1 | 8 | 19174035 | 19163904 | 14047513 | 12278119 |
| NEB-5 | NEBNext® Small RNA Library Prep | NEBNext | B | 2 | 8 | 15726206 | 15722621 | 11405023 | 9970595 |
| NEB-6 | NEBNext® Small RNA Library Prep | NEBNext | B | 3 | 8 | 16928701 | 16923441 | 11911815 | 10426736 |
| NEB-7 | NEBNext® Small RNA Library Prep | NEBNext | C | 1 | 8 | 16730535 | 16726418 | 12112546 | 10564774 |
| NEB-8 | NEBNext® Small RNA Library Prep | NEBNext | C | 2 | 8 | 15268930 | 15265435 | 11143421 | 9725521 |
| NEB-9 | NEBNext® Small RNA Library Prep | NEBNext | C | 3 | 8 | 14799992 | 14790986 | 10741517 | 9388746 |
| NEB-10 | NEBNext® Small RNA Library Prep | NEBNext | D | 1 | 8 | 17239669 | 17227653 | 12615888 | 11128918 |
| NEB-11 | NEBNext® Small RNA Library Prep | NEBNext | D | 2 | 8 | 15296072 | 15286892 | 11222041 | 9856961 |
| NEB-12 | NEBNext® Small RNA Library Prep | NEBNext | D | 3 | 8 | 13911293 | 13902739 | 10249997 | 9039781 |
| NEB-13 | NEBNext® Small RNA Library Prep | NEBNext | E | 1 | 8 | 14504103 | 14486624 | 10216851 | 8901109 |
| NEB-14 | NEBNext® Small RNA Library Prep | NEBNext | E | 2 | 8 | 16124825 | 16098301 | 11348331 | 9923236 |

| | | | | | | | | | |
|---|---|---|---|---|---|---|---|---|---|
| NEB-15 | NEBNext® Small RNA Library Prep | NEBNext | E | 3 | 8 | 17988105 | 17961291 | 12491862 | 10874180 |
| NEB-16 | NEBNext® Small RNA Library Prep | NEBNext | RA | 1 | 8 | 10668375 | 10434329 | 1612712 | 1069063 |
| NEB-17 | NEBNext® Small RNA Library Prep | NEBNext | RA | 2 | 8 | 10511673 | 10334495 | 1559628 | 1068297 |
| NEB-18 | NEBNext® Small RNA Library Prep | NEBNext | RA | 3 | 8 | 9929323 | 9873132 | 1535537 | 1047992 |
| NEB-19 | NEBNext® Small RNA Library Prep | NEBNext | HC | 1 | 8 | 9965698 | 9728438 | 1265436 | 864246 |
| NEB-20 | NEBNext® Small RNA Library Prep | NEBNext | HC | 2 | 8 | 8499421 | 8285821 | 1532582 | 1059428 |
| NEB-21 | NEBNext® Small RNA Library Prep | NEBNext | HC | 3 | 8 | 8928427 | 8603240 | 1667562 | 1144843 |

S2 Table

Intra-rater reliability of the three library preparation kit replicates of Mix A to Mix E and RA and healthy control samples measured by ICC(3,1) and absolute agreement

| Sample | ICC values | NEXTflex | NEBNext |
|---|---|---|---|
| Mix A | ICC | **0.995** | **0.993** |
| | lbound | 0.995 | 0.992 |
| | ubound | 0.996 | 0.994 |
| Mix B | ICC | **0.970** | **0.996** |
| | lbound | 0.967 | 0.995 |
| | ubound | 0.973 | 0.996 |
| Mix C | ICC | **0.995** | **0.996** |
| | lbound | 0.994 | 0.995 |
| | ubound | 0.995 | 0.996 |
| Mix D | ICC | **0.995** | **0.996** |
| | lbound | 0.995 | 0.996 |
| | ubound | 0.996 | 0.997 |
| Mix E | ICC | **0.994** | **0.934** |
| | lbound | 0.993 | 0.927 |
| | ubound | 0.995 | 0.941 |
| Rheumatoid arthritis patients | ICC | **0.991** | **0.992** |
| | lbound | 0.990 | 0.991 |
| | ubound | 0.992 | 0.993 |
| Healthy controls | ICC | **0.990** | **0.993** |
| | lbound | 0.989 | 0.992 |
| | ubound | 0.991 | 0.993 |

lbound = lower confidence interval bound; ubound = upper confidence interval bound

S3 Table

Pearson correlation coefficients of the read counts within the replicates of the kits for the synthetic miRNA samples

| | NEBNext-Mix.A.1 | NEBNext-Mix.A.2 | NEBNext-Mix.A.3 |
|---|---|---|---|
| NEBNext-Mix.A.1 | 1.000 | 0.992 | 0.995 |
| NEBNext-Mix.A.2 | 0.992 | 1.000 | 0.994 |
| NEBNext-Mix.A.3 | 0.995 | 0.994 | 1.000 |

| | NEXTflex-Mix.A.1 | NEXTflex-Mix.A.2 | NEXTflex-Mix.A.3 |
|---|---|---|---|
| NEXTflex-Mix.A.1 | 1.000 | 0.996 | 0.995 |
| NEXTflex-Mix.A.2 | 0.996 | 1.000 | 0.995 |
| NEXTflex-Mix.A.3 | 0.995 | 0.995 | 1.000 |

| | NEBNext-Mix.B.1 | NEBNext-Mix.B.2 | NEBNext-Mix.B.3 |
|---|---|---|---|
| NEBNext-Mix.B.1 | 1.000 | 0.996 | 0.995 |
| NEBNext-Mix.B.2 | 0.996 | 1.000 | 0.996 |
| NEBNext-Mix.B.3 | 0.995 | 0.996 | 1.000 |

| | NEXTflex-Mix.B.1 | NEXTflex-Mix.B.2 | NEXTflex-Mix.B.3 |
|---|---|---|---|
| NEXTflex-Mix.B.1 | 1.000 | 0.997 | 0.958 |
| NEXTflex-Mix.B.2 | 0.997 | 1.000 | 0.956 |
| NEXTflex-Mix.B.3 | 0.958 | 0.956 | 1.000 |

| | NEBNext-Mix.C.1 | NEBNext-Mix.C.2 | NEBNext-Mix.C.3 |
|---|---|---|---|
| NEBNext-Mix.C.1 | 1.000 | 0.996 | 0.995 |
| NEBNext-Mix.C.2 | 0.996 | 1.000 | 0.996 |
| NEBNext-Mix.C.3 | 0.995 | 0.996 | 1.000 |

| | NEXTflex-Mix.C.1 | NEXTflex-Mix.C.2 | NEXTflex-Mix.C.3 |
|---|---|---|---|
| NEXTflex-Mix.C.1 | 1.000 | 0.996 | 0.995 |
| NEXTflex-Mix.C.2 | 0.996 | 1.000 | 0.993 |
| NEXTflex-Mix.C.3 | 0.995 | 0.993 | 1.000 |

| | NEBNext-Mix.D.1 | NEBNext-Mix.D.2 | NEBNext-Mix.D.3 |
|---|---|---|---|
| NEBNext-Mix.D.1 | 1.000 | 0.996 | 0.996 |
| NEBNext-Mix.D.2 | 0.996 | 1.000 | 0.997 |
| NEBNext-Mix.D.3 | 0.996 | 0.997 | 1.000 |

| | NEXTflex-Mix.D.1 | NEXTflex-Mix.D.2 | NEXTflex-Mix.D.3 |
|---|---|---|---|
| NEXTflex-Mix.D.1 | 1.000 | 0.997 | 0.995 |
| NEXTflex-Mix.D.2 | 0.997 | 1.000 | 0.994 |
| NEXTflex-Mix.D.3 | 0.995 | 0.994 | 1.000 |

| | NEBNext-Mix.E.1 | NEBNext-Mix.E.2 | NEBNext-Mix.E.3 |
|---|---|---|---|
| NEBNext-Mix.E.1 | 1.000 | 0.913 | 0.912 |
| NEBNext-Mix.E.2 | 0.913 | 1.000 | 0.995 |
| NEBNext-Mix.E.3 | 0.912 | 0.995 | 1.000 |

| | NEXTflex-Mix.E.1 | NEXTflex-Mix.E.2 | NEXTflex-Mix.E.3 |
|---|---|---|---|
| NEXTflex-Mix.E.1 | 1.000 | 0.995 | 0.996 |
| NEXTflex-Mix.E.2 | 0.995 | 1.000 | 0.992 |
| NEXTflex-Mix.E.3 | 0.996 | 0.992 | 1.000 |

| | NEBNext-RA.1 | NEBNext-RA.2 | NEBNext-RA.3 |
|---|---|---|---|
| NEBNext-RA.1 | 1.000 | 0.993 | 0.993 |
| NEBNext-RA.2 | 0.993 | 1.000 | 0.991 |
| NEBNext-RA.3 | 0.993 | 0.991 | 1.000 |

| | NEXTflex-RA.1 | NEXTflex-RA.2 | NEXTflex-RA.3 |
|---|---|---|---|
| NEXTflex-RA.1 | 1.000 | 0.990 | 0.992 |
| NEXTflex-RA.2 | 0.990 | 1.000 | 0.990 |
| NEXTflex-RA.3 | 0.992 | 0.990 | 1.000 |

| | NEBNext-HC.1 | NEBNext-HC.2 | NEBNext-HC.3 |
|---|---|---|---|
| NEBNext-HC.1 | 1.000 | 0.993 | 0.992 |
| NEBNext-HC.2 | 0.993 | 1.000 | 0.993 |
| NEBNext-HC.3 | 0.992 | 0.993 | 1.000 |

| | NEXTflex-HC.1 | NEXTflex-HC.2 | NEXTflex-HC.3 |
|---|---|---|---|
| NEXTflex-HC.1 | 1.000 | 0.989 | 0.991 |
| NEXTflex-HC.2 | 0.989 | 1.000 | 0.989 |
| NEXTflex-HC.3 | 0.991 | 0.989 | 1.000 |

### S4 Table

Inter-rater reliability of the first replicate of each library preparation kit in Mix A to Mix E, RA and healthy control samples measured by ICC(3,1) and absolute agreement

| Sample | ICC | lBound | uBound |
|---|---|---|---|
| Mix A | 0.825 | 0.810 | 0.840 |
| Mix B | 0.825 | 0.810 | 0.840 |
| Mix C | 0.824 | 0.809 | 0.838 |
| Mix D | 0.818 | 0.803 | 0.833 |
| Mix E | 0.800 | 0.783 | 0.816 |
| Rheumatoid arthritis patient | 0.961 | 0.958 | 0.964 |
| Healthy controls | 0.955 | 0.951 | 0.959 |

lbound = lower confidence interval bound; ubound = upper confidence interval bound

S5 Table

Inter-rater correlation of the read counts within the first replicate of all five mixes in each kit for the synthetic miRNAs, rheumatoid arthritis (RA) and healthy control (HC) samples

| | NEXTflex -Mix.A.1 | srLp-Mix.A.1 | NEBNext-Mix.A.1 | QIAseq-Mix.A.1 | TailorMix -Mix.A.1 | CleanTag -Mix.A.1 |
|---|---|---|---|---|---|---|
| NEXTflex-Mix.A.1 | 1.000 | 0.842 | 0.869 | 0.789 | 0.830 | 0.819 |
| srLp-Mix.A.1 | 0.842 | 1.000 | 0.866 | 0.779 | 0.893 | 0.948 |
| NEBNextNext-Mix.A.1 | 0.869 | 0.866 | 1.000 | 0.764 | 0.794 | 0.852 |
| QIAseq-Mix.A.1 | 0.789 | 0.779 | 0.764 | 1.000 | 0.766 | 0.777 |
| TailorMix-Mix.A.1 | 0.830 | 0.893 | 0.794 | 0.766 | 1.000 | 0.902 |
| CleanTag-Mix.A.1 | 0.819 | 0.948 | 0.852 | 0.777 | 0.902 | 1.000 |

| | NEXTflex -Mix.B.1 | srLp-Mix.B.1 | NEBNext-Mix.B.1 | QIAseq-Mix.B.1 | TailorMix -Mix.B.1 | CleanTag -Mix.B.1 |
|---|---|---|---|---|---|---|
| NEXTflex-Mix.B.1 | 1.000 | 0.845 | 0.873 | 0.794 | 0.828 | 0.818 |
| srLp-Mix.B.1 | 0.845 | 1.000 | 0.871 | 0.786 | 0.880 | 0.950 |
| NEBNextNext-Mix.B.1 | 0.873 | 0.871 | 1.000 | 0.772 | 0.791 | 0.852 |
| QIAseq-Mix.B.1 | 0.794 | 0.786 | 0.772 | 1.000 | 0.762 | 0.774 |
| TailorMix-Mix.B.1 | 0.828 | 0.880 | 0.791 | 0.762 | 1.000 | 0.886 |
| CleanTag-Mix.B.1 | 0.818 | 0.950 | 0.852 | 0.774 | 0.886 | 1.000 |

| | NEXTflex -Mix.C.1 | srLp-Mix.C.1 | NEBNext-Mix.C.1 | QIAseq-Mix.C.1 | TailorMix -Mix.C.1 | CleanTag -Mix.C.1 |
|---|---|---|---|---|---|---|
| NEXTflex-Mix.C.1 | 1.000 | 0.841 | 0.869 | 0.784 | 0.827 | 0.805 |
| srLp-Mix.C.1 | 0.841 | 1.000 | 0.867 | 0.779 | 0.894 | 0.949 |
| NEBNextNext-Mix.C.1 | 0.869 | 0.867 | 1.000 | 0.767 | 0.800 | 0.851 |
| QIAseq-Mix.C.1 | 0.784 | 0.779 | 0.767 | 1.000 | 0.763 | 0.768 |
| TailorMix-Mix.C.1 | 0.827 | 0.894 | 0.800 | 0.763 | 1.000 | 0.892 |
| CleanTag-Mix.C.1 | 0.805 | 0.949 | 0.851 | 0.768 | 0.892 | 1.000 |

| | NEXTflex -Mix.D.1 | srLp-Mix.D.1 | NEBNext-Mix.D.1 | QIAseq-Mix.D.1 | TailorMix -Mix.D.1 | CleanTag -Mix.D.1 |
|---|---|---|---|---|---|---|
| NEXTflex-Mix.D.1 | 1.000 | 0.839 | 0.861 | 0.781 | 0.828 | 0.803 |
| srLp-Mix.D.1 | 0.839 | 1.000 | 0.866 | 0.775 | 0.892 | 0.952 |
| NEBNextNext-Mix.D.1 | 0.861 | 0.866 | 1.000 | 0.762 | 0.796 | 0.851 |
| QIAseq-Mix.D.1 | 0.781 | 0.775 | 0.762 | 1.000 | 0.765 | 0.762 |
| TailorMix-Mix.D.1 | 0.828 | 0.892 | 0.796 | 0.765 | 1.000 | 0.873 |
| CleanTag-Mix.D.1 | 0.803 | 0.952 | 0.851 | 0.762 | 0.873 | 1.000 |

| | NEXTflex -Mix.E.1 | srLp-Mix.E.1 | NEBNext-Mix.E.1 | QIAseq-Mix.E.1 | TailorMix -Mix.E.1 | CleanTag -Mix.E.1 |
|---|---|---|---|---|---|---|
| NEXTflex-Mix.E.1 | 1.000 | 0.849 | 0.813 | 0.793 | 0.823 | 0.816 |
| srLp-Mix.E.1 | 0.849 | 1.000 | 0.802 | 0.785 | 0.889 | 0.949 |
| NEBNextNext-Mix.E.1 | 0.813 | 0.802 | 1.000 | 0.732 | 0.746 | 0.786 |
| QIAseq-Mix.E.1 | 0.793 | 0.785 | 0.732 | 1.000 | 0.772 | 0.776 |
| TailorMix-Mix.E.1 | 0.823 | 0.889 | 0.746 | 0.772 | 1.000 | 0.878 |
| CleanTag-Mix.E.1 | 0.816 | 0.949 | 0.786 | 0.776 | 0.878 | 1.000 |

| | NEXTflex-Mix.RA.1 | srLp-Mix.RA.1 | NEBNext-Mix.RA.1 | QIAseq-Mix.RA.1 | TailorMix -Mix.RA.1 | CleanTag -Mix.RA.1 |
|---|---|---|---|---|---|---|
| NEXTflex-Mix.RA.1 | 1.000 | 0.966 | 0.966 | 0.962 | 0.968 | 0.961 |
| srLp-Mix.RA.1 | 0.966 | 1.000 | 0.960 | 0.953 | 0.979 | 0.983 |
| NEBNext-Mix.RA.1 | 0.966 | 0.960 | 1.000 | 0.943 | 0.957 | 0.954 |
| QIAseq-Mix.RA.1 | 0.962 | 0.953 | 0.943 | 1.000 | 0.964 | 0.939 |
| TailorMix-Mix.RA.1 | 0.968 | 0.979 | 0.957 | 0.964 | 1.000 | 0.970 |
| CleanTag-Mix.RA.1 | 0.961 | 0.983 | 0.954 | 0.939 | 0.970 | 1.000 |

| | NEXTflex-Mix.HC.1 | srLp-Mix.HC.1 | NEBNext-Mix.HC.1 | QIAseq-Mix.HC.1 | TailorMix -Mix.HC.1 | CleanTag-Mix.HC.1 |
|---|---|---|---|---|---|---|
| NEXTflex-Mix.HC.1 | 1.000 | 0.968 | 0.964 | 0.962 | 0.968 | 0.948 |
| srLp-Mix.HC.1 | 0.968 | 1.000 | 0.958 | 0.955 | 0.979 | 0.964 |
| NEBNext-Mix.HC.1 | 0.964 | 0.958 | 1.000 | 0.940 | 0.954 | 0.945 |
| QIAseq-Mix.HC.1 | 0.962 | 0.955 | 0.940 | 1.000 | 0.961 | 0.924 |
| TailorMix-Mix.HC.1 | 0.968 | 0.979 | 0.954 | 0.961 | 1.000 | 0.954 |
| CleanTag-Mix.HC.1 | 0.948 | 0.964 | 0.945 | 0.924 | 0.954 | 1.000 |

S6 Table

Inter-rater correlations between rlog read counts and the theoretical sample concentration of the 40 non-equimolar miRNA oligonucleotides

| Kit | Mix | Replicate | Pearson correlation |
|---|---|---|---|
| NEXTflex | A | 1 | 0.59 |
| NEXTflex | A | 2 | 0.58 |
| NEXTflex | A | 3 | 0.57 |
| NEBNext | A | 1 | 0.40 |
| NEBNext | A | 2 | 0.40 |
| NEBNext | A | 3 | 0.42 |
| NEXTflex | B | 1 | 0.31 |
| NEXTflex | B | 2 | 0.32 |
| NEXTflex | B | 3 | 0.22 |
| NEBNext | B | 1 | 0.31 |
| NEBNext | B | 2 | 0.31 |
| NEBNext | B | 3 | 0.32 |
| NEXTflex | C | 1 | 0.48 |
| NEXTflex | C | 2 | 0.47 |
| NEXTflex | C | 3 | 0.48 |
| NEBNext | C | 1 | 0.33 |
| NEBNext | C | 2 | 0.34 |
| NEBNext | C | 3 | 0.33 |
| NEXTflex | D | 1 | 0.25 |
| NEXTflex | D | 2 | 0.24 |
| NEXTflex | D | 3 | 0.24 |
| NEBNext | D | 1 | 0.24 |
| NEBNext | D | 2 | 0.26 |
| NEBNext | D | 3 | 0.25 |
| NEXTflex | E | 1 | 0.58 |
| NEXTflex | E | 2 | 0.58 |
| NEXTflex | E | 3 | 0.58 |
| NEBNext | E | 1 | 0.41 |
| NEBNext | E | 2 | 0.40 |
| NEBNext | E | 3 | 0.40 |
